# FLOCK STASIS DRIVES FLYING SPEED IN PIGEONS, WHILE ARTIFICIAL MASS ADDITIONS DO NOT

**DOI:** 10.1101/2022.09.24.509309

**Authors:** Daniel W. E. Sankey, Steven J. Portugal

## Abstract

Animals are characterised, in part, by their use of voluntary movement, which is used to explore and exploit resources from their surrounding environment. Movement can therefore benefit animals, but will cost them their energetic reserves. Thus, adaptations for faster movements with negligible increases in energy expenditure will likely evolve via natural selection. Individual and social-level mechanisms have been shown to optimise this speed/energetic trade-off. Nevertheless, studies of social-level traits typically ignore individual variation, which is a cornerstone principle in evolutionary ecology. Furthermore, how individual phenotype interacts with the phenotypic composition of the group to govern the cost of transport may have been entirely overlooked. We investigate speed and the energetic consequences of individual-level phenotypic differences using body mass (both natural and artificially manipulated with additional weights) of homing pigeons (*Columba livia*) (*N* =16 birds; *N* = 193 useable flight trajectories). We then turn to social level phenomena, and manipulate the composition of pigeon groups by body mass (*N*= 12 birds in four treatments; *N* = 192 useable flight trajectories) and leadership rank (*N* = 30 birds in three groups, *N* = 286 useable flight trajectories) following earlier leadership identification flights (*N* = 33 birds, *N* = 306 useable flight trajectories). “Natural” body mass was predictive of flying speed in solo flights, but not in groups of greater mass by composition; “artificial” mass loading had no impact on speed in solo fliers, and was not tested in groups. Groups of *leader phenotypes*, showed faster speeds, and greater cohesion than *follower phenotype* groups, both in terms of flock spread, but also in consistency of positioning within the flock (“flock stasis”) across the flight. Flock stasis was further analysed across all other group flights. Its positive impact on speed was found to be consistent across all experimental treatments. Therefore, predicting flock stasis may be critical to understanding optimal phenotypic compositions of birds, and thus the social evolution of birds which fly together. We provide evidence that greater stasis may be driven by phenotypic compositions (i.e. groups of leaders, and homogeneous mass groups) and also discuss the implications of stasis for different flocking structures (e.g. V-formations) and human crowd control.

## INTRODUCTION

The evolution of voluntary movement is not unique to animals, but is a defining characteristic [1] that likely evolved to allow exploration and exploitation of the environment [2, 3]. Expanding the range or speed of movements may provide more foraging opportunities through increasing the rate of resource acquisition and by reaching farther away resources [4, 5], though this may also come at a greater energetic cost [6, 7]. Optimal foraging theory predicts that individuals that can move faster will increase their success of resource acquisition via an increased encounter rate, while assuming faster speeds come at a greater energetic cost [5, 8]. This assumption might not always hold true as both individual and social-level strategies can offset the costs of movement [9–14].

At the individual level, physical adaptations can reduce the cost of travel. For example, body streamlining can reduce drag (and thus energetic cost) in aquatic [9, 10] and aerial vertebrates [12, 15]. Also, individual-level behavioural strategies may have evolved to optimise travel through the optimal use of their “energy landscape”. For example, birds that make use of predictable rising air (e.g. thermals – columns of rising air) [14, 16], or adaptive combinations of active and passive dispersal through water currents in juvenile fish [17].

At the social-level, behavioural interactions with conspecifics have evolved to optimise travel costs. For example, in V-formation flights, vortices produced from the wingtips of leading individuals [18, 19] may explain decreased energetic output of trailing individuals [20], when relative body position and the timing of their flaps are spatially and temporally “in-phase” [19]. Similarly, fish have been shown to decrease muscle activity in artificial vortices [21]. However, current studies of social-level energy-saving strategies still implicitly ignore individual variation, which is a foundational principle in evolutionary ecology [22]. Furthermore, for social species, an individual’s success is intrinsically tied not only to its own phenotype, but also to the phenotypic composition of its group [23]. Overall, how individual phenotype and group phenotypic composition interact to govern the speed and costs of travel is an important but unknown aspect of the life history of animals which travel in groups.

Homing pigeons (*Columba livia*) are well suited to answer questions about speed and the costs of movement, and the interaction of individual-level phenotype and group phenotypic composition for a number of reasons. Pigeons need to use flight (a costly form of locomotion [24]) to navigate home, and can do so alone [25–27], or in groups [28–30]. Group composition and group size are easy to manipulate in pigeons – by releasing birds in groups of predetermined phenotype. Pigeons also exhibit high robustness to the application of animal attached biologgers, which can be used to measure speed (using GPS; e.g. [27, 31]) and energetic proxies (using accelerometers; e.g. [32, 33]). Through the application of biologging technology, measures of morphological and behavioural phenotypes are easily attainable in pigeons, including repeatable “in-flight” phenotyping (which is rare in biologging studies REF). For example, repeatable measures of speed (solo and group) and leadership (group) are easily measured with GPS loggers, due to the reliability of their homing [27–29], and the consistent transient leadership hierarchies shown to be stable in pigeon flocks [28–30,34,35]. Biologging also has less welfare impact on captive animals, due to the relatively short-term attachment (<1 day; [36]), and the provision of surplus food to restore their energy levels after deployment. Finally, the extensive historical study of pigeons provides a wealth of baseline knowledge, for more proficient predictions and inference of results.

Previously, we found an example of how energetic costs may relate to individual phenotype and the phenotypic composition of pigeon groups. In birds, energetic output may be optimised (minimised) by flying at an individually specific optimum speed, which we found may be characterised by their morphological phenotype. Flying any faster than this optimum will require additional energy output, through increased chemical energy required to power flight muscles; as will flying slower, via a reduction in lift produced by momentum [37]. Birds were found to compromise from their preferred speed to fly with the group, which highlights that the benefits of grouping (e.g. enhanced predator avoidance [38, 39] or decision making accuracy [26, 40]) may outweigh these costly compromises. Heavy and light pigeons – which are thought to optimise their energetic profile by flying faster and slower respectively, may have to increase their relative energetic output to fly together. Therefore, an individual of a given mass (individual phenotype) would benefit energetically by flying in groups of similar mass (group phenotypic composition) as the group may fly closer to the individual’s preferred speed, which would require less *energetically costly* compromise. This highlights the potential of individual-level and social-level behavioural and morphological traits that may result in more effective travel. In this paper, we explore how individual-level phenotype and group phenotypic compositions may interact to govern speed and the energetic costs of bird flight, via experimental manipulations, to provide a novel synthesis of these domains.

### INDIVIDUAL-LEVEL PHENOTYPE: BODY MASS AND ARTIFICIAL MASS MANIPULATIONS

It is widely demonstrated that morphological phenotype has a close relationship to speed and the costs of flight [6,37,41]. In particular, mass and wing-loading have been shown to be predictive of speed in birds across species [6]. Intra-species differences in body mass have also been shown to predict greater flight speeds in pigeons [27]. Furthermore, faster travel in powered flight is thought to be associated with greater energetic costs in birds [37, 42], fitting the predictions of optimal foraging theory [5, 8]. Heavier birds will require a faster air flow to produce the same amount of lift [37]. A concurrent increase in energetic output is expected for heavier birds, because of the higher weight-per-area for each flap (wing loading) [37]. Nevertheless, this has not been tested, and it is possible that heavier individuals could potentially mitigate the costs of their faster speeds through efficient flight kinematics, as has been shown for birds carrying additional mass loads [43, 44]. For example, cockatiels (*Nyphicus hollandicus*) have been shown to slow down while retaining their flapping frequency [44]. Alternatively, zebra finches (*Taeniopygia guttata*) have been shown to increase flapping and decrease wing amplitude in response to artificial mass [43]. However this latter study enforced a speed of 10m/s [43], and therefore the zebra finches were not capable of responding by slowing down. Here, we ask whether larger birds naturally employ some of these same tactics to mitigate the costs of faster speeds, and second whether we can induce these same tactics by manipulating the mass of individuals artificially.

### SOCIAL-LEVEL BEHAVIOURAL TRAITS: FLOCK LEADERSHIP MANIPULATIONS

Studies of collective movement suggest that consistent behavioural differences may also play a role in inter-group differences in optimising the trade-off between speed and energetic cost. Usherwood et al. [45] found that flying in a pigeon flock comes at an energetic cost over flying solo, with denser flocks exhibiting greater flapping frequencies, even after accounting for speed. Therefore, any method which can manipulate (decrease) the density of bird flocks could provide a method to reduce energetic cost without concurrent reductions in speed. Johnstone and Manica [46] provide a potential solution using an evolutionary model of leadership dynamics. They demonstrated that populations of leader phenotypes – owing to their goal-oriented nature – suffer reduced coordination/cohesion as a result of attempting to dominate movement decisions [46]. Together, these studies suggest that groups of individuals with *on-average* greater leadership could form less dense flocks [46], and further, owing to this reduction in density, will fly further/faster per unit energy [45]. Here, we ask the question as to whether a behavioural phenotype (leadership) can manifest in more effective flight when alterations are made to the group phenotypic composition of the flock.

### SOCIAL-LEVEL MORPHOLOGICAL TRAITS: GROUP MASS COMPOSITION MANIPULATIONS

Given that heavier birds fly faster both across [6] and within species([27], manipulations of the group composition by body mass provides a way to test the impact of group phenotypic composition on the relationship between speed and energetic costs. As mentioned above, individuals of a similar mass may also prefer to fly at a similar speed and homogeneous mass group compositions may reduce speed compromise [47], which may be highly costly in birds [37]. Such a reduction would highlight a benefit individuals can achieve at the social level, via optimal choice of group membership [23]. The resulting energetic benefits for individuals could therefore be specific to their own phenotype, and the phenotypic composition of their group [23]. Nevertheless, this question remains unstudied at present. Here we ask whether manipulations of the *body mass* composition can influence speed and energetic proxies across the social context (i.e. phenotypic composition).

### ALTERNATIVE SOCIAL-LEVEL STRATEGIES: FLOCK STASIS

Inferring benefit from more efficient use of speed and/or energetic expenditure in less dense flocks (see *Social level behavioural traits*, above) is problematic, as birds which form dense clusters will do so for a reason. Whether the function of dense cluster-formations is of navigational benefit [26,33,40] or anti-predator benefit [38, 39] is not approached in this study, but are both of potential importance for the fitness of individuals in groups [48]. Therefore we expanded our search for optimal strategies beyond *flock density*, and into *flock stability* (or, stasis). Specifically, by flock stasis, we are referring to the potential for differing degrees of consistency in the spatial positioning in some flocks over others, which may be based upon their phenotypic composition. Flying in a flock has been shown to increase energetic expenditure of birds over flying solo [33], which is thought, in part, to be attributable to the unpredictability and/or inconsistency (i.e. heterogeneity) of airflows left in the wake of birds in front [33, 45]. Stasis in flock positioning could make the air environment more predictable and/or homogenous. We expect individuals of a similar mass to exhibit greater stasis, due to their comparatively lower levels of speed compromise, which may manifest in accelerations (by lighter, slower birds) or decelerations (by heavier, larger birds), which retain cohesion but may modify the air environment negatively compared with birds which fly at a consistent speed. Given the expected stability of the air environment, we expected higher stasis to be predictive of flock speed. Additionally, we expect that groups of leader phenotypes will exhibit lower stasis than groups of follower phenotypes, due to a reduction of “attempted leadership initiations” in the follower phenotypes [46, 49]. Therefore how group phenotypic composition influences flock stasis, and whether flock stasis is predictive of optimisation of the speed to energetic cost trade-off was another question approached in the present study.

### METHODOLOGY AND PREDICTIONS

We experimentally manipulated individual mass (artificially), the group leadership composition, and the group body-mass composition of homing pigeons to test *a priori* predictions (in brackets) through experimental manipulations as follows in three separate experiments. Furthermore we conducted a comparative analysis of all group flights to investigate flock stasis (described above). **In experiment one**, by artificially manipulating the mass of pigeons, we predicted (1) reductions in speed while energetic costs remain stable – following the results of experiments which allowed speed changes in response to artificial mass loadings [44]. We also predict (2) that body mass will positively covary with speed and energetic proxies regardless of treatment. **In experiment two**, by predetermining leadership hierarchies, we first separated birds into groups of leaders and followers. We predicted that (3) groups of leaders would show reduced density (following [46]), and with this, (4) reduced energy expenditure (following [45]). **In experiment three**, if body mass predicted solo speed in individuals, we predicted (5) that group flights comprising more heavy individuals would exhibit greater flock speeds. We also predicted (6) that heterogeneous groups (i.e. those with a mixture of heavy and light individuals) would experience greater costs of flight relative to flying in homogenous groups, as groups will need to reach a speed consensus – via costly speed compromise – to avoid splitting. We then combined all group flights for **our comparative analysis** to look for evidence of flock stasis, which we predicted would (7) enhance flock speed and/or decrease energetic proxies. Additionally we predicted greater stasis in (8) homogeneous body-mass groups over heterogeneous compositions, and (9) groups of “followers” over groups of “leaders”.

## METHODS

Homing pigeons (*N* = 49) were kept in purpose-built lofts at Royal Holloway University of London (Surrey, UK; latitude = 51.416, longitude = -0.572), and provided with food (Johnstone & Jeff Four Season Pigeon Corn, Gilberdyke, U.K.), water, and grit (Versele-Laga - Colombine Grit and Redstone, Deinze, Belgium) *ad libitum* throughout the course of the study period (April—September 2018).

### Experiments and general protocol

The birds were weighed (CoffeeHit: Coffee Gear Digital Bench Scale – 2kg/0.1g limit/accuracy) weekly, providing a key morphological covariate (“natural” bird mass), while simultaneously monitoring welfare [50]. Repeatability of body mass was deduced using likelihood ratio tests, with 95% confidence intervals estimated using 10,000 parametric bootstrap iterations.

The study comprised three experimental flocks; (1) *Artificial mass manipulations of birds flying solo* (*N* = 16 birds, age = 2.5 years old, dates of study: April—May 2018, with previous experience from the release site; see below), (2) *Group leadership composition manipulations* (*N* = 33 birds, age = 9 months, dates of study: July—September 2018, with no experience of the release site at the beginning of the study), and (3) *Group body mass composition manipulations* (*N* = 12 birds, assorted by body mass, from the same batch as in leadership composition manipulations, dates of study: September 2018).

Our key parameters of interest across the experiments were flight speed (m/s), and energetic proxies *i)* flap frequency (Hz), and *ii*) dorsal body amplitude (mm). To record these variables we adopted a biologging approach, using GPS (speed; 5Hz, QStarz BT-Q1300ST, Düsseldorf, Germany; mass = 12.5g) and accelerometers (flap frequency and dorsal body amplitude; 200Hz; AX3, Axivity Ltd, Newcastle upon Tyne, UK; 8g). See *SI text* for further information on logger treatment and attachment.

Each flight experiment comprised releasing the homing pigeons from a site between Windsor Castle and Eton (coordinates: latitude = 51.497, longitude = -0.589) away from their home lofts at Royal Holloway (coordinates: Latitude = 51.415, Longitude = -0.573). Firstly, birds were gathered from their home loft and placed into wicker carrying baskets (dimensions = 80cm x 40cm x 22cm) for transportation to release site. Following this, the birds were driven 8.90km north (exact bearing = - 0.07rad) to the release site. The birds were typically in transit for 15-25 minutes (traffic depending). Older birds (experiment one) had experience from the site, for training protocol on younger birds (experiments two and three) see *SI text*. At the release site, both GPS and the accelerometers were switched on and attached to the back of each bird. GPS loggers were switched on at least five minutes before deployment to ensure an accurate signal was being received from the satellites. The birds were released either *i*) solo (experiment one), in the order they were randomly selected from the box, and following a minimum of 10 minutes delay from the release of the previous bird, or *ii*) as groups (experiments two and three) by opening the side hatch on the wicker basket following at least a 15 minute delay from the release of a previous group. This time-delay ensured birds were not subsequently meeting up en-route from the release site to the home lofts.

### Experiment one: solo flight mass manipulations

For experiment one (solo mass manipulations), the purpose was to manipulate the mass of individuals to determine the impact of this mass manipulation on solo flight speed [27] and flight kinematics (*sensu* [43, 44]). Individuals either flew without any mass manipulation, as a control condition (∼0g, Velcro strip), or with 5g or 10g additional mass added to their backs using tin wheel-balancing bicycle weights (Abba ME, Essex, UK). The self-adhesive weights were attached to the accelerometer loggers (on the cranial end of the pigeons’ backs) to ensure the additional artificial mass was close to the centre of the pigeon’s body mass, which is located towards the head [51].

Given the initial mass of the birds (range 409.6g – 533.1g), the mass load as a proportion of body mass (logger mass plus additional mass load by bird mass) ranged from 3.8% for heaviest birds in the 0g artificial mass treatment, to 7.4% for the lightest birds in the 10g artificial mass treatment. This means that a commonly cited 5% rule was surpassed in some cases in the present study [36]; however as mentioned, impact of biologging may be mitigated in captive animals relative to wild species (see *Introduction* and [36]). Additionally, we were able to provide data on the response of pigeons to the additional mass, which could inform ethical decision making in future work.

Each flight release event contained a mix of the three mass loading treatments (i.e. control, 5g and 10g), to regulate for temporal differences in weather conditions – which may perturb speed, despite us controlling for wind conditions in the analyses (see *Speed, Tortuosity and Wind parameters*, below; also see *SI text* and Table S1 for randomisation procedures). Before the termination of the study, we explored the data for any evidence of pigeons “pairing up” (see *Computational methods* below), to, firstly, remove the data from the analysis, and second, to ensure that each individual had completed at least three solo flights per mass condition. Determination of whether birds flown under solo treatments did subsequently pair up was defined as: less than 50m from a neighbour (following [27]) for over 50% of the flight was stated as being a pair. Following removal of paired flights, the number of solo flights per mass condition were 0g (N = 60 flights); 5g (N = 66 flights); 10g (N = 67 flights) (see Table S4).

### Experiment two: group leadership composition manipulations

For experiment two (group leadership composition manipulations), we firstly randomly allocated 33 pigeons into three separate groups. Following a training phase (see *SI text* for details on training), each of the three pigeon groups (N = 11, N = 11, N = 10 for each group respectively, following one loss in training flights) were released ten times from the standard Windsor-Eton release site to establish leadership hierarchies (subsequently referred to as “leadership identification flights”).

Following the allocation of leadership ranks (following [29]; also see below), each group was subdivided into leaders (top 5 leadership scores) and followers (bottom 5 scores). In the groups of 11 (*N* = 2 groups) the individual with the middle score (at leadership rank 6) was left-out of further study. The new groups (*N* = 6 groups; three groups of leaders and three groups of followers) were released a further 10-11 times (Table S3) to assess differences in group dynamics between groups of leaders and groups of followers (subsequently referred to as “leadership manipulation flights”). Bird losses (i.e. birds which did not return from a particular flight; *N =* 3), as well as all logger failures (N = 6), and flights where one or more groups did not participate. A maximum of one flight was missed per group, due to individuals undergoing routine procedures; all flights are documented in Table S3. Groups were not changed as a result of losses but instead group size (which was thus diminished in some cases) was treated as a categorical variable with fixed effects in our models (see *Statistics*).

### Experiment three: group mass composition manipulations

For experiment three (“mass manipulation flights”), we took 27 pigeons with body mass distribution approximately normal (Shapiro Wilks test; *W* = 0.962, *p* = 0.392), and formed two subsequent groups: ‘light’ and ‘heavy’. Six light individuals and six heavy individuals (randomly selected from the bottom/top eight of the 27 birds, respectively) were selected, leaving a difference of 46.9 g between the heaviest bird from the light group (mean = 374.3 g; S.D. = 19.3 g) and lightest bird from the heavy group (mean = 455.3 g; S.D. = 15.4 g). On a given flight/day the groups were either flown as complete but separate heavy/light groups (homogeneous mass groups), or with two individuals swapped into the other group (heterogeneous mass groups) before flight. See *SI text* and Table S2 for randomisation of group compositions. Therefore we utilised four distinct mass compositions, 1) “all light” (N = 6 ‘light’ birds), 2) “predominantly light” (N = 4 ‘light’ birds; N = 2 ‘heavy’ birds), 3) “predominantly heavy” (N = 4 ‘heavy’ birds; N = 2 ‘light’ birds), and 4) “all heavy” (N = 6 ‘heavy’ birds). We flew each composition eight times, totalling 192 trajectories, with no missing data or birds. We express these conditions as the proportion of heavy individuals in the flock, therefore we had eight group flights with each [0], [0.33], [0.67] and [1], as a proportion heavy birds [square brackets are used to denote the proportion of heavy birds in this study throughout].

### Computational methods

#### Fission, cohesion and sensitivity algorithms

We removed all GPS timestamps for the first and last 1000m of the flights (following [32]) to compare only relatively steady sections of the return flight home. Further, in group flights, we removed data where the distance of an individual was over 50m to the centroid (mean of latitude and longitude across group at each timestamp; see *SI text*). We also ran statistics on centroid distances of 25m and 75m to test the sensitivity of the statistics to our arbitrary choices of parameters. Fission – the proportion of flight separated from the group – was then calculated by dividing the total number of timestamps removed by the total number of steps.

We only recorded further metrics for each flight if a group remained stable in their composition. This was defined as *unchanged group membership* for a proportion of over 0.1 (10%) of the flight (subsequently referred to as “minimum flight proportion”). This allowed us to reduce erroneous readings, which may be caused by different group sizes or different group compositions, which come about by fission and/or fusion of birds. However, it was therefore possible to record multiple readings of a single metric in one flight (i.e. a reading for each of the different group compositions which remained stable for over 10% of the flight). Therefore we recorded the date and unique flight ID, to use as random intercepts (see *Statistics*), to deal with pseudoreplication [52]. We tested the sensitivity our subsequent analyses to our arbitrary choice of “minimum flight proportion”, by testing “minimum flight proportion” values of 0.05 and 0.25, as well as 0.1 (see *SI text*).

#### Leadership

To assess leadership, we used pairwise correlations analysis on the merged trajectories (see methods in [29]), whereby time-lags of similar movements are used to quantify the directional correlation delay of their turns (and hence the leadership). For example, if one individual turns (on average) 0.2s before the rest of the flock, it will be considered a leader; whereas another individual which turns 0.2s after the rest of the flock, it will be considered a follower. Leadership analysis was not considered if individuals did not remain cohesive (<50m; following [27]) for over 50% of the flight. If the resulting “leadership” matrix from all flights demonstrates significant transitive hierarchy (following [27– 30,34,53]), we can separate flocks by leadership rank to test our hypotheses (i.e. to test the relationship between leadership, flock composition and flock density and the efficiency of travel; see *Predictions*).

#### Speed, tortuosity, and wind variables

We measured ground speed using point-to-point distance travelled between GPS coordinates at each time-step (5Hz), for both individuals and groups (“flock speed”, using centroid coordinates). Tortuosity – which is a key component to control for in models of speed [54] – was measured by the difference in heading between each time-step. We took a median of speed and tortuosity for the entire flight (solo flights), or for sections of flight with different, but stable, group compositions (group flights; see above). The efficiency of each flight was classically calculated as the perfect beeline (straight line between the site and the loft) divided by the total distance travelled [26, 55]. This metric was calculated for each individual/group for each flight.

As speed can be influenced by wind speed and direction relative to the direction of travel [54], we record wind speed (accuracy ± 0.1 m/s) and direction (accuracy ± 22.5 degrees) every 0.5 hours at the home loft using a weather station (Aercus Instruments WS4083 Pro Wireless Weather Station; Greenfrog Scientific, Doncaster, U.K). Using the nearest timestamp of wind data to the first timestamp of trimmed GPS pigeon data, we calculated wind currents parallel (support wind) and perpendicular (cross wind) to direction of travel (following [54]). A mean of support and cross wind components were calculated across the whole flight (solo), or for periods of flight with stable group composition as above (group).

#### Energetic proxies: accelerometer measures

Data were downloaded from the accelerometry loggers as .svg files, and then processed to .csv files in OMGUI (https://github.com/digitalinteraction/openmovement/wiki/AX3-GUI), before exporting to R [56]. We then trimmed accelerometer data to match the start and end of the timestamps from the trimmed GPS (see above). Following this, we calculated flap frequency and dorsal body amplitude for each pigeon, for each flight. Flapping frequency was calculated using smoothed dorsal acceleration (z-axis of accelerometer; smoothed over 0.025s), and removing static acceleration (over 15 wingbeat cycles, >2s, see [32]), before estimating upper reversal point (see [19]) as a measure of each flap cycle. This measure was calculated at each wingbeat (i.e. time between one flap and the next, in flaps per second, Hz). The amplitude (mm) of the dorsal body acceleration (“dorsal body amplitude”) was estimated via double integration of the acceleration curve, before passing through a Butterworth filter with cut-off frequency 2.5Hz (see [32, 45] for further information). Again, this measure was calculated at each wingbeat. Medians of accelerometer measures were taken per bird, per flight, in all experimental treatments.

#### Flock density and stasis

In group flights, we calculated a further series of covariates describing the flock density. Across the flock, all neighbour-to-neighbour distances (in metres) were collated and averaged (mean average) at each time-step distance (Fig. 1). Flock spread was defined as the median value of these per-time-step values across sections of flight where group composition remained stable (as in speed, tortuosity and wind parameters; see above). Dorso-ventral and cranio-caudal spread were defined as the distance (in metres) between the furthest individuals to the left/right or front/back, respectively, with regard to the heading of the centroid. Consistent with flock spread this was recorded at each time-step, and then further reduced to a median value across cohesive sections of flight to provide measures of dorso-ventral spread and cranio-caudal spread for further analysis.

**Figure 1.**
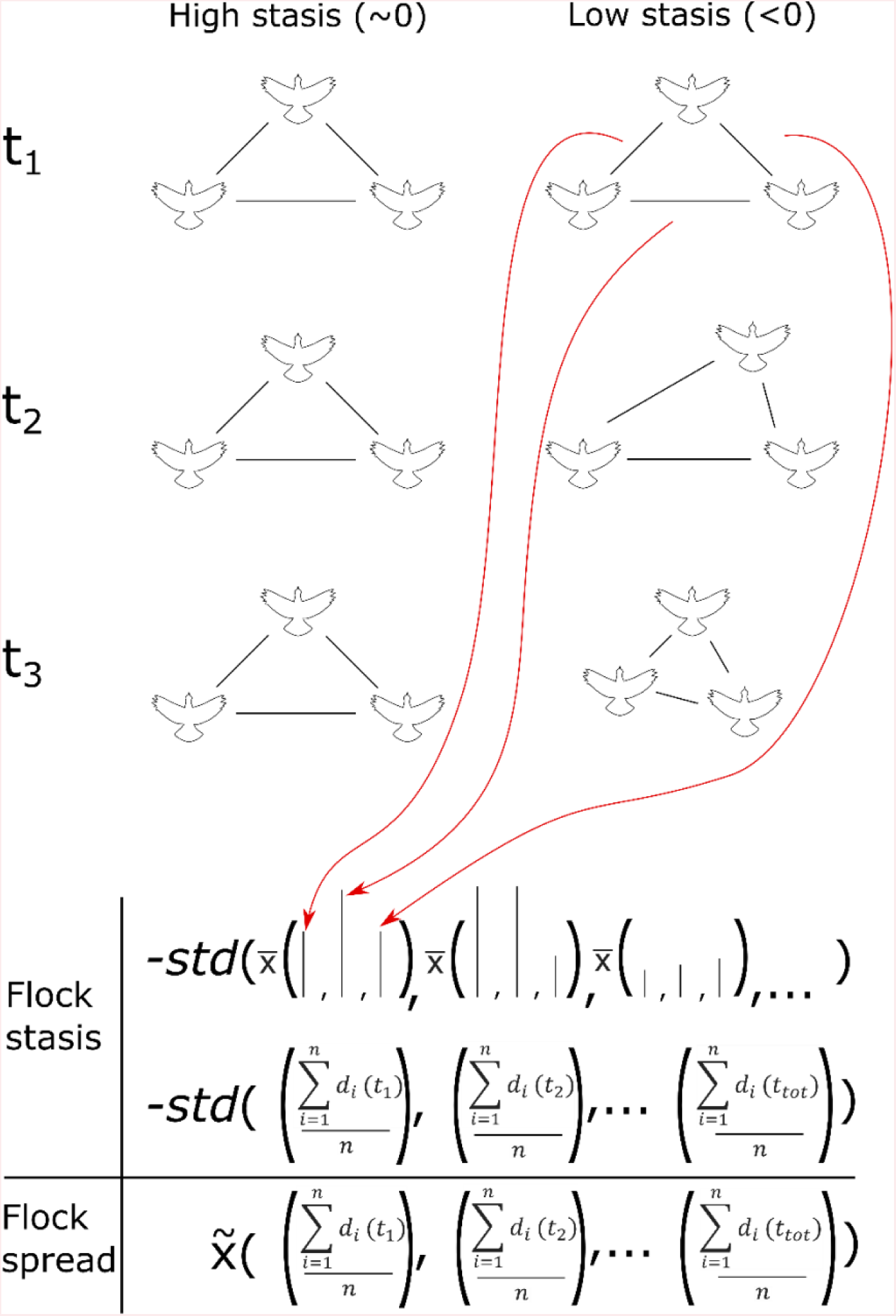
Flock stasis and flock spread calculation schematic. Distances (m) between each individual (black lines) are calculated between each bird and for each time-step (t_1_, t_2_, etc.). These are then compiled (red arrows), summed and divided by the total number of distances (mean average, 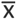) at each time-step. Our measure of flock stasis is the inverse of the standard deviation (*std*) of these data (units = -*std*(m)). Our measure of flock spread is a median value (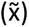) of the same data (units = m). Birds in the left column represent a highly static flock, with values for flock stasis close to zero; birds in the right column represent a flock with low (more negative) stasis, due to individual’s moving around more.

Additionally, following our rationale and predictions for social-level strategies which may confer optimal use of the aerodynamic environment, we measured flock stasis, a novel measure which estimates the variability in inter-individual positioning across the flock. Flock stasis (a measure of variation in flock spread) was calculated in using flock spread data, but using the *inverse of the standard deviation* instead of a *median* of the data from all time-steps (Fig. 1). Therefore we present flock stasis with units (-*std*(m)), reflecting the input variable (distance, m) and the statistic (standard deviation, *std*) used to derive the measure.

### Statistics

We used linear mixed models (R package “nlme” [57]) to estimate the explanatory power of various covariates on dependent variables – solo speed, flock speed, flap frequency, dorsal body amplitude, and flocking parameters (e.g. flock stasis; above). Fixed effects were as follows: experimental treatment (i.e. artificial mass load – solo flights, and group composition – group flights); group size (after fission and bird losses; see above); support-wind and cross-wind components; “natural” – non-manipulated – bird mass was a fixed variable in solo flights, and tested via group composition changes in group flights; finally, flocking parameters were used as fixed effects when they were not treated as dependent variables (e.g. in models of flock speed). Our random effects included: pigeon ID (in solo flights and group flights without group level measures); group number (in experiment two; as the same treatment was conducted across three separate groups); unique flight ID and date (unique flight ID and date picking up smaller and larger scale local perturbations in temporal environmental conditions); finally, when we compared all data from experiments two and three (cf. our comparative analysis; see *Predictions*); experimental treatment was also included as a random intercept (i.e. “leadership Identification flights”, “leadership manipulation flights”, and “mass manipulation flights”). All model fits were tested to for a fit to the assumption of parametric residuals, and, depending on the greatest visual coherence of model residuals with a fitted qq-norm plot (REF)m were treated with either *i*) no transformation, *ii*) Box-Cox transformations [58], or *iii*) log transformations.

In our model estimating flock speed, collinearity of flocking parameters (e.g. flock spread, flock stasis) was dealt with by removal of less statistically significant parameters over a threshold value of correlation coefficients (|*r*|>0.5), following [59] (see Fig S1 for a schematic and detailed description of this process).

## RESULTS

### Experiment 1: Solo flights with artificial mass manipulations

#### Artificial mass manipulations

Following prediction (1), that artificially mass manipulated birds would slow down, while keeping energetic proxies – flap frequency and dorsal body amplitude – constant was not supported. There was no evidence that manipulating mass loading (5g weights, 10g weights) decreased ground speed in pigeons (N = 16 birds; maximum N = 15 flights per bird: five flights in each of the three conditions) either as a numeric variable (LME; DF = 64, *t* = -0.766, *p* = 0.447) or a categorical variable (Fig. 2*A*; LME with Tukey’s pairwise post-hoc test; DF = 63; comparisons: 0g - 5g – *t.ratio* = 0.380, *p* = 0.924; 0g - 10g – *t.ratio* = 0.772, *p* = 0.721; 5g - 10g – *t.ratio* = -0.425, *p* = 0.905; see Table S4 for all statistics). Additionally, flap frequency did not remain stable, it showed significant increases when artificial mass load was treated as a numeric variable (Fig. 2*B*; LME; DF = 64, *t* = 2.067, *p* = 0.0428), however, not when treated as categorical variables (Table S3). The increase in flap frequency was only slight, with an increase of approximately 0.02Hz per gram of added artificial mass. Dorsal body amplitude also changed across mass load, the observed amplitude was significantly lower when treated as a numeric variable (LME; DF = 64, *t* = -4.682, *p* < 0.001) and across categorical group (10g condition) when compared with 0g and 5g treatments (Fig. 2*C*; LME with Tukey’s pairwise post-hoc test; DF = 63; comparisons: 0g - 10g – *t.ratio* = 4.760, *p* < 0.001; 5g - 10g – *t.ratio* = 3.712, *p* = 0.001). Decreases in amplitude were more marked than differences in flap frequency, with an estimated decrease of approximately 0.17mm per additional gram of artificial mass. In additional analyses, we found that support wind component and cross wind components had a significant impact on ground speed (positive and negative respectively; support wind – *t* = 2.680, *p* = 0.009; cross wind – *t* = -3.432, *p* = 0.001).

**Figure 2.**
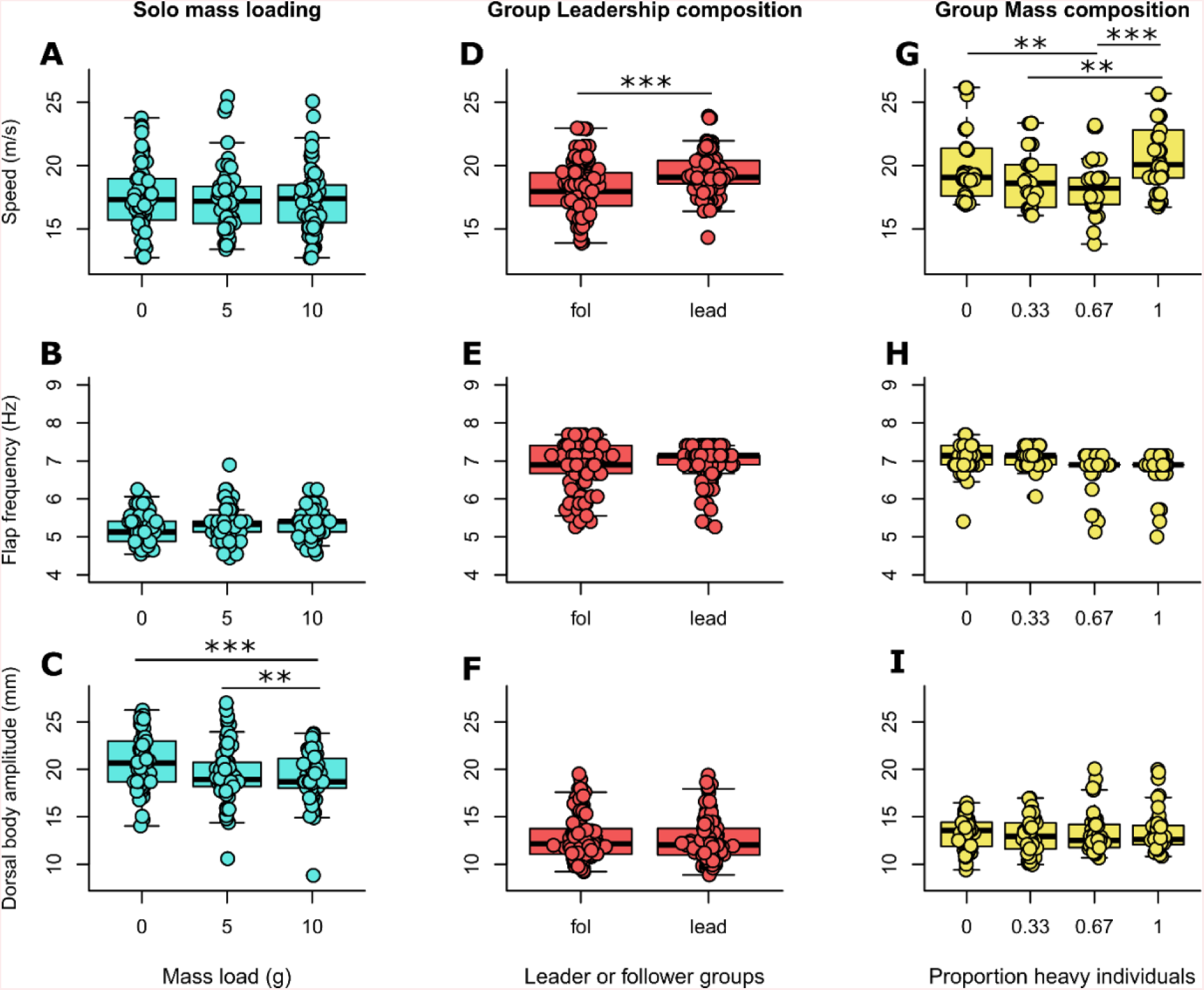
Individual level flight metrics from all three experiments. Median flight speeds (m/s; top row), flapping frequencies (Hz; middle row) and dorsal body acceleration values (mm; bottom row) are given as points with box and whisker plots for each experiment as follows: **(A-C)** Solo mass loading (turquoise): flights with additional artificial mass loadings attached to the accelerometer logger either 0g (Velcro strip), 5g or 10g (see *Methods*). **(D-F)** Group leadership composition (orange-red): metrics for individuals in groups of followers (fol) or leaders (lead) (see *Methods*). **(G-I)** Group mass composition (yellow): group flights with different proportions of heavy individuals; either 0 (six light individuals), 0.33 (four light and two heavy individuals), 0.67 (two light and four heavy individuals), or 1 (six heavy individuals). Statistically significantly different responses between groups are provided from a Tukey posthoc test of a linear mixed effects model (see *Methods*), where p value is either < 0.05 (*), < 0.01 (**), or < 0.001 (***).

#### Solo speeds and accelerometer metrics

As expected from prediction (2), “natural” (non-manipulated) bird body mass was a predictor of flight speed, regardless of artificial mass treatment, with an estimated increase in speed of 0.1 m/s per gram of additional “natural” body mass (Fig. 3*A*; LME; DF = 168, *t* = 5.564, *p* < 0.001). As predicted, body mass was also associated with higher flap frequencies (Fig. 3*B*; LME; DF = 167, *t* = 3.255, *p* = 0.001), and lower dorsal body amplitudes (Fig. 3*C*; DF = 167, *t* = -2.503, *p* = 0.013). Individual measurements of “natural” bird mass were also highly repeatable – repeatability analysis, using likelihood ratio tests, with 95% confidence intervals estimated using 10,000 parametric bootstrap iterations; *R* = 0.80 ± 0.03 (s.e.m.), 95% CI: 0.73–0.85, *p* < 0.001.

**Figure 3.**
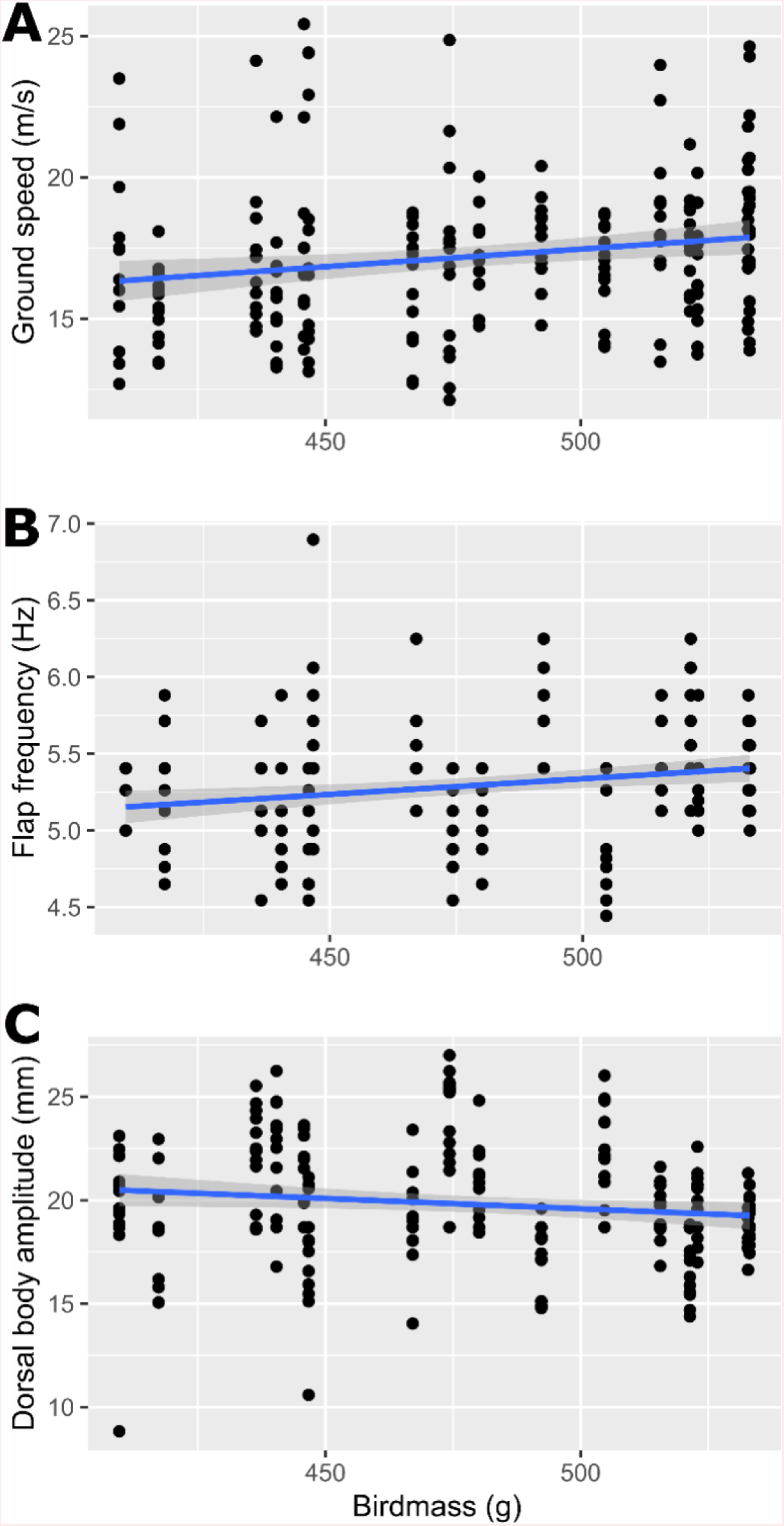
The effect of “natural” bird mass on speed and energetic metrics. Bird mass (g) (x-axis) is plotted (points) with fitted linear model (blue line with grey shaded 95% confidence intervals, fitted in ggplot [60]), against **A)** Median ground speed (m/s), **B)** Median flap frequency (Hz), and **C)** Dorsal body amplitude (mm). All relationships were statistically significant – see above in *Results: Solo flights – speeds and accelerometer metrics*.

### Experiment 2: group leadership composition manipulations

#### Leadership identification

All three groups exhibited significantly transitive leadership hierarchies (methods from [28]; *T_1_* = 1.000, *T_1_* = 1.000, *T_1_* = 0.890; *p* < 0.001 for all groups), and so were divided into groups with the five lowest and five highest leadership scores (directional correlation delay; see *Methods*) per group. Body mass was not predictive of leadership score in any of the three groups (LM; DF = 8; Group 1: *t* = -0.494, *p* = 0.635; Group 2: *t* = 0.334, *p* = 0.747; Group 3: *t* = 2.254, *p* = 0.054), nor was there any significant difference in the body mass of leaders and followers (mean mass of followers/leaders = 403.03g/413.16g respectively; T-test; *t* = 0.841, DF = 26.624, *p* = 0.408), thus suggesting body mass was not a significant factor in determining leadership.

#### Effect of group phenotypic composition

Contrary to our expectations in prediction (3), leader flocks had increased density in all measures of density, with decreased flock spread (LME with negative Box-Cox transformation; DF = 15, *t* = 1.975, *p* = 0.067), decreased cranio-caudal spread (LME with negative Box-Cox transformation; DF = 15, *t* = 0.743, *p* = 0.469), but only significant decreases observed in dorso-ventral spread (LME with negative Box-Cox transformation; DF = 15, *t* = 2.376, *p* = 0.031). Further analyses of dorso-ventral spread found this to be the *flock-density* measure with the highest predictability of flock speed (see models from our comparative analysis below). Therefore, dorso-ventral spread is arguably the most important density measure we recorded, given the major thesis of our present work (cf. speed and the costs of transport).

Prediction (4) was also not supported, as the leader flocks, which were more dense, demonstrated greater speeds than follower flocks (Fig. 2*D*; LME; DF = 28, *t* = -6.087, *p* < 0.001), with an estimated speed increase of approximately 1.19m/s for leader groups, from 18.11m/s (± 1.98 (S.D)) to 19.30m/s (± 1.53 (S.D)). Flock speeds for training flights were closer to the speed of followers – 18.13m/s (± 1.53 (S.D.). We found no effect of leadership composition on energetic proxies flap frequency (LME; DF = 27, *t* = 0.9607, *p* = 0.345) or dorsal body amplitude (LME; DF = 27, *t* = 0.212, *p* = 0.833).

In additional analyses, we found that leader groups exhibited less fission than follower groups, though not significantly less (mean 19% and 12% of flights spent separated from the group in follower and leader groups respectively; LME with negative Box-Cox transformation; DF = 50, *t* = 1.562, *p* = 0.125). There were no significant differences in route efficiency between leader and follower groups (LME; DF = 50, *t* = 1.200, *p* = 0.236), with follower group mean efficiencies at approximately 0.73, and leaders at 0.79 (where the efficiency of a beeline is 1).

### Experiment three: group mass composition manipulations

We found no support for prediction (5); that flocks with a greater number of heavy individuals by proportion would fly faster. When treated as a numeric variable, the proportion of heavy birds was not predictive of flight speed (LME; DF = 136, *t* = 1.000, *p* = 0.319). However we did find that certain group mass compositions were predictive of speed over others (Fig. 3*G*; LME with Tukey’s pairwise post-hoc test; DF = 134; comparisons, given as proportion of heavy individuals in group [in square brackets] are as follows: [0 - 0.67] – *t.ratio* = 3.287, *p* = 0.007; [0.33 – 1] – *t.ratio* = -3.341, *p* = 0.006; [0.67 – 1] – *t.ratio* = -4.402, *p* < 0.001, see Table S4 for all statistics). The most significant differences were observed between homogeneous groups ([0] and [1]) and heterogeneous groups ([0.33] and [0.67]) where the homogeneous mass groups flew an estimated 1.56m/s faster (using mean values).

We also found no support for prediction (6), that energetic proxies (flap frequency and dorsal body amplitude) would be reduced in groups of homogeneous composition. Flap frequency was greater, and dorsal body amplitude was lower in homogenous groups, but not significantly so in either case (flap frequency: LME; DF = 28, *t* = 0.396, *p* = 0.694; dorsal body amplitude: LME; DF = 28, *t* = 0.396, *p* = 0.480), When treated as a categorical or numeric variable (Fig. 3*H,I*; see Table S4 for statistics), the closest value to significance for energetic proxies across groups was flap frequency between [0] and [0.67] tukey corrected p value DF = 161, t.ratio – 2.353, tukey corrected p-value = 0.090. For dorsal body amplitude, despite heavier groups [0.67] and [1] showing some of the highest values (20.05mm and 19.98mm), the medians were lower (13.56mm, 12.94mm for lighter groups [0] and [0.33] respectively; vs. 12.50mm and 12.60mm for heavier groups [0.67] and [1] respectively), and there was no difference across the groups (greatest difference was between [0.33] and [1]; DF = 161, t.ratio = - 0.409, *p*= 0.974).

In additional analyses, fission was not found to be different across groups of differing mass composition. The mean fission for each mass-composition group as follows: [0] = 7%, [0.33] = 9%, [0.67] = 11%, [1] = 15% (see Table S7 for all statistics). There were no differences in route efficiency between groups of different mass composition, with mean route efficiency for each group as follows: [0] = 0.95, [0.33] = 0.94, [0.67] = 0.92, [1] = 0.95 (see statistics in Table S7).

### Comparative analysis: density, stasis and speed across group-flight treatments

#### Flock speed models

In support of prediction (7), the selected model predicting flock speed (See *Methods* and Fig. S1) across all group flight experiments, showed that both flock stasis and density (dorso-ventral spread), as additive covariates (Table 1) had significant impact on group flight speed (Table 1; Fig. 4; LME; DF = 100, Flock stasis – *t* = 5.590, *p* < 0.001; Dorso-ventral spread *t* = -2.760, *p* = 0.007; for diagnostics see Fig. S2). Flock stasis, as a predictor, was robust to a sensitivity analysis on chosen parameters “Fission distance” and “Minimum flight proportion” (see *Methods* and Table S6), and as a predictor of flight speed across each experiment (Table 1). Simultaneously there were no significant changes in the mean group flap frequency or mean dorsal body amplitude in more static flock flights. This was only performed on the experimental conditions (“leadership manipulation flights”, and “mass manipulation flights”), however, and not the “leadership identification flights”, as the accelerometer data were not analysed due to time constraints (flap frequency: LME; DF = 48; *t* = 0.576, *p* = 0.567; dorsal body amplitude: LME; DF = 48, *t* =0.176, *p* = 0.861). Experimental treatment “Leadership composition” as a factor, was still a significant predictor of speed in a model with flock stasis added (LME; DF = 14, *t* = 3.96, *p* = 0.001). Conversely, group mass composition was not predictive of speed in models including flock stasis (homogeneous vs. heterogeneous: LME; DF = 14, *t* = -1.130, *p* = 0.27). This implies that flock stasis does not fully describe the variation in group speed in leader vs. follower flocks but may describe the differences in the groups of different mass composition, i.e. whether homogeneous or not.

**Figure 4.**
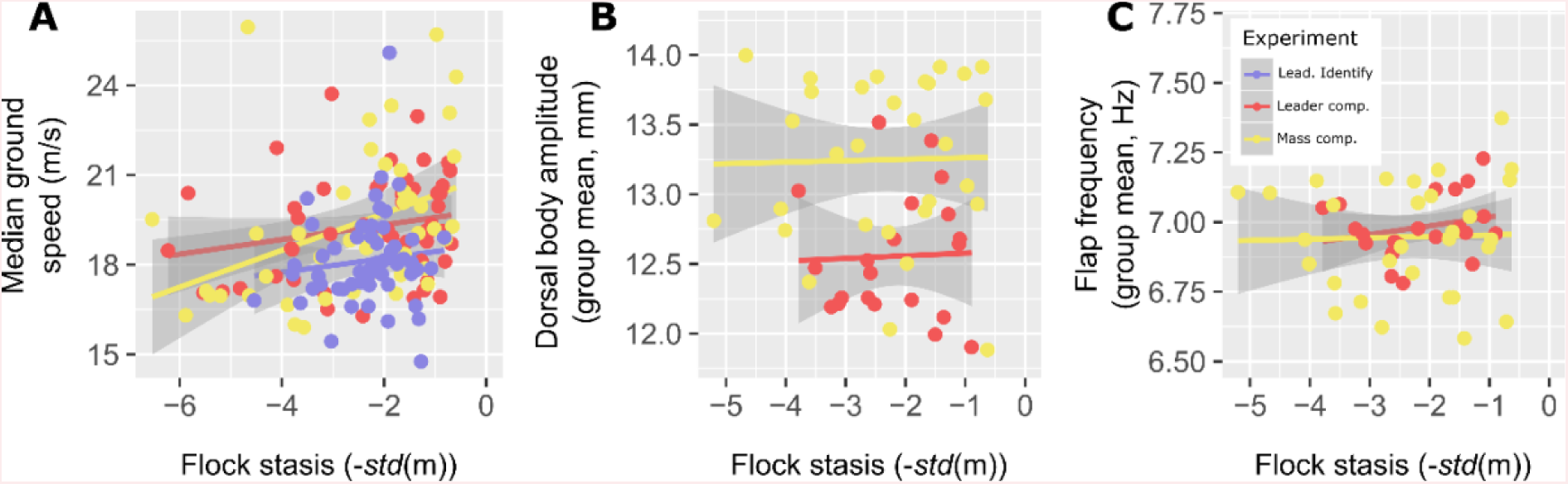
Flock stasis is predictive of greater speed, without increases in energetic proxies. Flock stasis (-*std*(m)) is plotted against **(A)** Median flock speeds (ground speed (m/s); points) for sections of flight where group composition remained stable (more than 10 % of a flight), **B)** Dorsal body amplitude (mm; mean of values across whole group/flight) and **C)** Flap frequency (Hz, mean values across whole group/flight). Linear models with 95% confidence intervals are fitted using R-package “ggplot2” [60]. Colours of lines and points correspond to leadership identification flights (purple), leadership composition manipulations (orange-red) and mass composition manipulations (yellow) (see legend in **(C)**). Accelerometer data used in the generation of **(B)** and **(C**) were not analysed for leadership identification flights.

**Table 1.** Selected model for flock speed. P-values (if applicable) are reported for the impact of flocking variables (“Flock stasis” and “Dorso-ventral spread”) on speed, and the model with the lowest AIC score (“Best model”, either one flocking variable “1var”, both flocking variables in an additive model “sum” or as an interaction “int”; See detailed methods in *SI text* and Fig. S1). Values are reported for each series of flights as well as a combination of all flights (columns: leadership composition manipulations – “Lead comp.”, mass composition manipulations “Mass comp.”, leadership identification flights “Lead. Identify”, and all flights combined “All”).

|  | **Lead comp.** | **Mass comp.** | **Lead. Identify** | **All** |
| --- | --- | --- | --- | --- |
| *Flock stasis* | 0.0002 | 0.0453 | 0.0353 | <0.001 |
| *Dorso-ventral spread* | NA | NA | NA | 0.007 |
| ***Best model*** | “1var” | “1var” | “1var” | “sum” |

In additional analyses, the impact of group size, when treated as a numerical covariate, was close to significant as a predictor of speed, with a negative relationship between the two variables (LME with negative Box-Cox transformation; DF = 111, *t* = 1.807, *p* = 0.074), it is therefore unlikely that the larger group sizes in leader flocks (due to greater losses in follower groups, see Table S3) explain the speed differences between the groups.

#### Flock stasis and flock density models

Contrary to prediction (8) groups of leader phenotype, had positive influence on flock stasis in group leadership manipulation experiments (Fig. 5*A*; Table S6; LME; DF = 15, *t* = 2.237, *p* = 0.041; this result held for 4 out of 5 trials testing for sensitivity of parameters “Fission distance” and “Minimum flight proportion”; See Table S6). Prediction (9), otherwise, was supported, as homogeneity of mass was predictive of flock stasis (Fig.5*A*; LME; DF = 14, *t* = -2.261, *p* = 0.040; homogeneity of masses held as significant in 3/5 of the sensitivity iterations).

**Figure 5.**
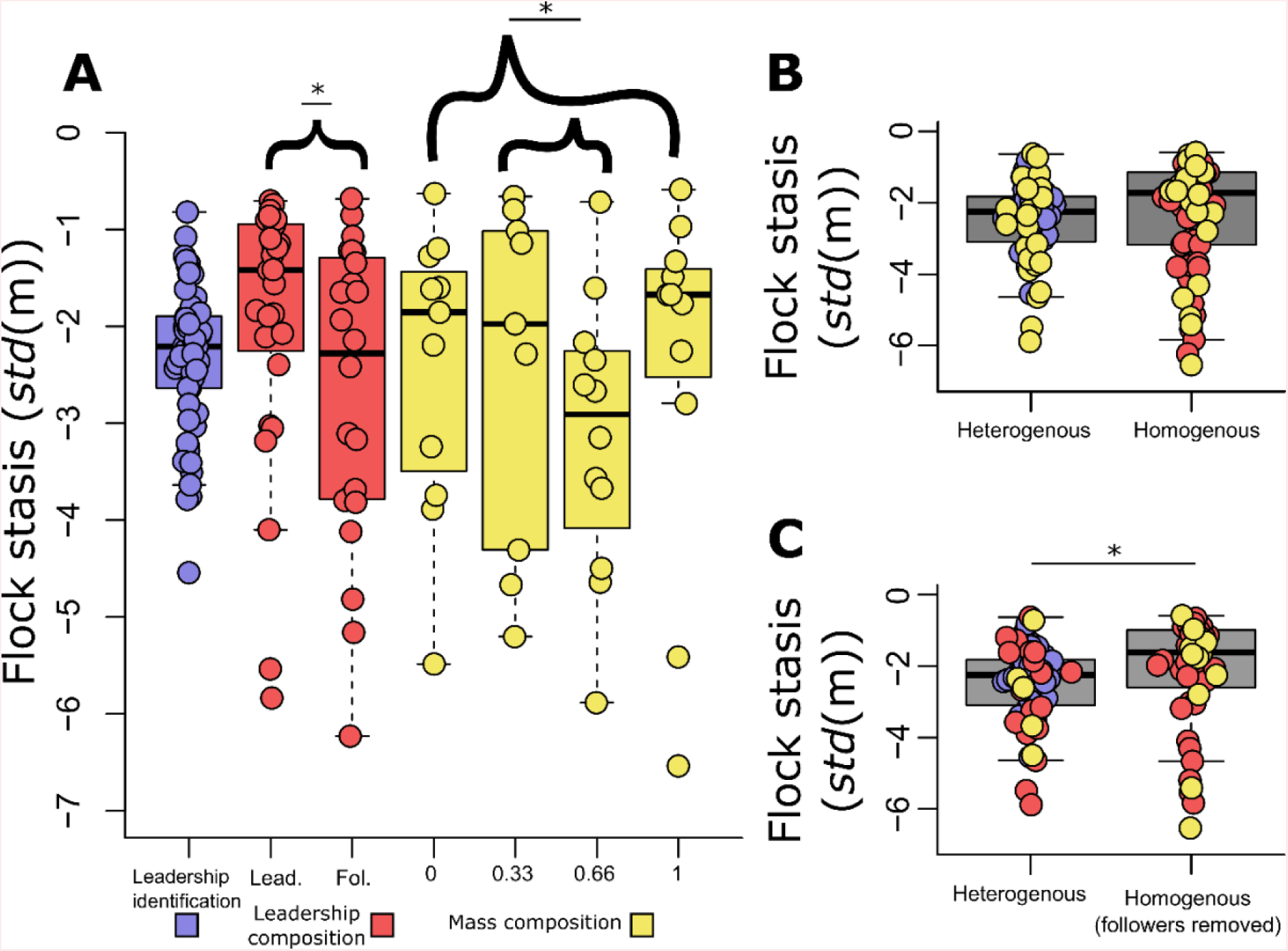
Flock stasis for groups of different phenotypic composition. Flock stasis (-*std*(m)) is plotted (points; box and whisker plots), against phenotypic composition. **A)** (From left to right) leadership identification flights (N=10-110), groups of leaders (N=4-5; Table S3), groups of followers (N=3-5; Table S3), mass manipulated groups (N=6) with proportions of heavy individuals either 0, 0.33, 0.67, 1 respectively. Statistically significantly different responses between groups are drawn for p-values of either < 0.05 (*), < 0.01 (**), or < 0.001 (***). The curly brackets above the mass manipulation experiments represent a statistical comparison between homogeneous groups (0 and 1 proportions of heavy individuals) and heterogeneous groups (0.33 and 0.67 proportion of heavy individuals). **B-C)** show a comparison of homogeneous groups and heterogeneous groups across all experiments, where heterogeneous groups are defined as: heterogeneous mass groups, and leadership identification flights; and homogeneous groups are defined as homogeneous mass groups, leader groups and follower groups, **B)** or with follower groups omitted **C)**. Colours correspond to leadership identification flights (purple), leadership composition manipulations (orange-red) and mass composition manipulations (yellow) (see colours in boxes below **A)**)

In additional analyses, homogeneous flocks, across all experiments, showed greater flock stasis (where leader flocks, follower flocks, and similar sized mass groups were treated as homogeneous; and leadership identification flights, and mixed mass groups were considered as heterogeneous). However this increase was not statistically significant unless flights from follower groups were taken out (Fig. 5; with followers included: Fig. 5*B*; LME; DF = 35, *t* = 1.649, *p* = 0.108; with followers excluded: Fig. 5*C*; LME; DF = 35, *t* = 2.080, *p* = 0.045; sensitivity analyses showed statistical significance for 1/5 iterations with followers included and 4/5 without; Table S6). Additionally, flight order was found to have negative impact on flock stasis, which was close to significance (LME; DF = 103, *t* = -1.927, *p* = 0.057), meaning more experienced flocks exhibited less stability in their positioning. Flock size, after accounting for individuals which split from the group, had a negative, but non-significant, relationship with flock stasis (LME; DF = 111, *t* = -1.388, *p* = 0.168).

Dorso-ventral spread (m), as reported above, was decreased in leader flocks compared with follower flocks (LME; DF = 15, *t* = 2.376, *p* = 0.031; the result held for 3/5 of the sensitivity analysis iterations – see Table S6), from a mean lateral flock spread of 5.12m in followers to 4.71m in leaders. This is despite leader flocks having a biased starting size due to greater pigeon losses in follower groups (Table S3), and flock size having a strong positive impact on dorso-ventral spread (LME with negative Box-Cox transformation; DF =110, *t* = -9.939, *p* < 0.001). Mass compositions were not predictive of dorso-ventral spread, in any condition, nor was homogeneity of flock composition across experiments (see Table S6). Flight order had a negative impact on dorso-ventral spread, indicating that groups were less compact in later flights, however the effect was not significant (DF = 102, *t* = 1.585, *p* = 0.116).

## DISCUSSION

We have identified mechanisms which can facilitate speed increases in groups. Specifically, groups which can remain highly static in their positioning (i.e. high flock stasis) appear to travel faster at no extra energetic cost, even when accounting for differences in the tortuosity of flight movements [54]. This suggests that group compositions which manage to keep stable flock structures can benefit energetically, while also observed to be flying in a denser cluster (i.e. on the dorso-ventral axis), which may confer additional anti-predator benefits [61, 62]. Although the most highly cited function of flying in groups is that of energetic saving [33], cluster-flocking actually has been shown to impose additional energetic cost over solo flight [33], especially in tighter clusters [33, 45]. Flock stasis may function to mitigate these additional costs. Such costs are thought to be produced by *i*) the increased control over movements to avoid collisions [33, 45], and *ii*) the production of inconsistent airflow which is likely to be a heterogeneous mix of upwash and downwash currents, which may be highly unpredictable [45]. Consistency in the positioning of individuals will likely increase either (or both) of the above, as the predictability of neighbours’ movements may reduce the risk of collisions (increasing the confidence of movement decisions [63]), and also the predictability and general homogeneity in the airflow left in the wake of other pigeons. Thus allowing birds to fly faster, and in more tightly compact groups, without necessitating concurrent increases in energetic output [33, 45], which will likely be adaptive when energetic constraints represent a substantial selective pressure [5,14,64].

We suggest that certain configurations of group phenotypic composition are predictive of high levels of stasis in flight, and furthermore that greater stasis allows faster flying speed. Nevertheless, a valid alternative hypothesis is that group phenotypic composition governs speed, and speed governs stasis. However, if phenotypic composition primarily governed speed, we would expect groups with a greater proportion of heavy individuals to fly faster, as individual mass and solo speed were highly correlated in experiment one [27]. This was not the case, instead, we observed that homogeneous mass groups travel faster than heterogeneous mass groups regardless of group mass by proportion. This suggests homogeneity in mass composition facilitated speed increases through increases in flock stasis. (See more discussion of group mass composition below, in section subtitled: *Social-level morphological traits: group mass composition manipulations*).

### INDIVIDUAL-LEVEL PHENOTYPE: BODY MASS AND ARTIFICIAL MASS MANIPULATIONS

While increasing mass by group composition was not indicative of flight speed, “natural” body mass of the birds was, and this held regardless of artificial mass load treatment. However, body mass – while predictive of speed, showed conflicting results in terms of energetic output, with greater flapping frequencies and lower dorsal body amplitude in heavier individuals. Nevertheless, despite this conflict, the energetic cost of steady, horizontal flight is thought to be proportional to the cube of flap frequency, while only to the square of wingbeat amplitude [65, 66]. Additionally, empirical work has suggested that wingbeats with higher frequency, low amplitude are necessary for (more costly) powered turning flight [45]. Therefore it is likely that heavier individuals are likely to pay a higher cost for their faster speeds, via their increases in flap frequency.

In artificially mass manipulated flight, our observed decrease in dorsal body amplitude had a greater effect size than the observed increase in flapping frequency. However it is highly unlikely that pigeons *save energy* while maintaining speed in artificially mass manipulated flight. To use an analogy of airplanes, one would be unlikely to power an Airbus A380 (560,000kg maximum take-off weight [67]) on the engine of an Airbus A320 (68,000kg maximum take-off weight [68]) [69]. Instead, it seems, as demonstrated theoretically [66] and empirically [45], increases in flap frequency may be much more costly than decreases in wingbeat amplitude, and our work further supports this point. Overall, we suggest that pigeons are robust to the application of moderate mass loadings (3.8%–7.4% of total body mass) because they have the capacity to adjust flight kinematics to retain their usual flying speed. This suggests either that *i*) speed is under high selection pressure [64], or that *ii*) the result of slowing down would reduce momentum induced lift to unsustainable levels [37].

Further increases in the mass load might reveal a critical point at which speed reduces, though this may result stop flight altogether, and would be better investigated in a wind tunnel due to ethical considerations [70]. Further *decreases* in logger mass might, on the other hand, prove more useful in free-flying pigeons for estimation of both *i*) the impact of biologgers, and *ii*) natural unloaded movement kinematics of birds [71]. As explained by Wilson et al. [71], kinematic data from a whole range of different device loadings can be, firstly, fitted with a regression model. Second, the impact of different mass loadings (slope) and the inferred “true” value of the kinematic metric when unloaded with biologgers (y-intercept) can be extrapolated from the model. (See [71] for an example plot with acceleration on the y-axis, device mass on the x axis, and a fitted linear regression line.)

We suggested that heavier individuals may mitigate the additional energetic costs of faster speeds, and that a similar flight kinematic response could be seen in mass loaded birds (albeit without the concurrent increases in speed). Indeed they increased their flap frequency and decreased their dorsal body amplitude (same for “artificially” and “naturally” heavier birds), but also increased speed (which the “artificially” loaded individuals did not). This suggests that the speed increases concurrent with additional mass is more than just a result of increased wing loading [6]. The pectoralis muscle is thought to be the major driver of wing suppression and the supracoracoideus muscle with elevation [64]. Our work suggests that the aforementioned muscles could be larger in heavier pigeons, with a concurrent increase (or “scaling up”) of the metabolic supply of chemical energy necessary to power these relatively larger muscles [64, 72].

### SOCIAL-LEVEL BEHAVIOURAL TRAITS: FLOCK LEADERSHIP MANIPULATIONS

We found that “leader groups” demonstrated higher density and flock stasis than “follower groups”, despite our predictions to the contrary. We initially reasoned that leaders would attempt more initiations [46, 73], and that this in turn would predict reduced density and stasis. However, our predictions were not supported, with leaders exhibiting significantly greater cohesion, and stasis, without an obvious decrease in energetic expenditure. As our birds were all flown from the same site, this may have reduced the conflict in navigational decision making [26, 48]. As leaders have been shown to learn routes better than followers (see [27]), this conflict may have actually *reduced* in leader flocks which all know the route better than follower flocks. Therefore, we suggest that with reduced conflict in route direction, the “leader groups” were able to *optimise their flock dynamics*, rather than *attempt initiations* toward their preferred direction of travel.

Leader pigeons may also exhibit a more goal-oriented individual-level phenotype [55, 74]. Pigeons with greater “peak-fidelity” – a measure of their coherence in their solo flight routes – predicted leadership in pigeon pairs [55]. Goal-orientedness was also shown to have a positive impact on leadership in fish, however only when balanced with moderate levels of social tendency, as fish which were highly goal-oriented would split from the group, reducing their influence [75]. As goal-orientedness is thought to represent a trade-off between propensity to risk isolation and the safety of the group [76], such a behaviour may correlate with individual differences in boldness [76, 77]. Indeed boldness has been found to predict leadership propensity in fish [77] and pigeons [74]. Additionally, faster pigeons have been shown to lead movements [27]. This might explain why “leadership composition” as a factorial variable had predictive power on flock speed in models – above the predictive power from the stasis variable alone. Altogether, disentangling the impact of route learning, route conflict, speed and goal-orientedness on flock stasis is a challenge for future research.

Our methods were designed to decrease flock density, and hence modify the speed/energetic trade-off, by forming groups with high conflict through increases in their natural “goal-orientedness” (see *Introduction* and [46]). This may have not been successful due to aforementioned *decreases* in conflict through enhanced route learning. Future work should potentially *i*) fly groups from different sites, or *ii*) fly the leaders of one group with the leaders from another group. Both these concerns were taken into consideration in the present study: firstly, our employed comparative analyses required consistency in the experimental set-up; second, by chance alone, individuals with intrinsic traits that confer a “leader phenotype” could be unevenly distributed across experimental groups, and mixing across groups could introduce unaccountable bias.

### SOCIAL-LEVEL MORPHOLOGICAL TRAITS: GROUP MASS COMPOSITION MANIPULATIONS

Increased flock stasis in homogeneous mass groups was expected, nevertheless our results suggest a different mechanism than the one proposed. We reasoned that, (1) in groups of a similar mass, we would find an intrinsic reduction in speed compromise between the birds. Further, (2) acceleration/deceleration responses would be less common (or indeed necessary), and thus, via moving around less, would result in greater levels of stasis. Following our results, this now seems unlikely, as groups of heavy birds did not fly faster than groups of slower birds, so speed compromise may not be the governing force we expected it to be. Instead we hypothesise the following: that *i*) the general morphological profile of individuals (e.g. wingspan, structural size; which has been shown to correlate with body mass in birds; [6, 78], will affect the flow rate and magnitude of air currents left in their wake and *ii*) individuals can achieve a more optimal trade-off between downwash avoidance and upwash exploitation in the wake of similar sized individuals. This is highly speculative, as no present evidence can confirm or reject this claim. However, a recent study suggests that a previous dichotomy between cluster-flocking and V-formation flight mechanisms [79] might not be as straightforward as first thought, with observations of “compound V” flock shapes within the cluster formations of shorebirds [80]. We know that flying in a flock comes at a cost in pigeons [33, 45], but this does not rule out the possibility for *relative* savings when individuals are matched in size and/or gait.

### COMPARATIVE ANALYSES

To attempt to identify the mechanism underpinning how some flocks are able to reach higher stasis, we also investigated whether homogeneous groups, more broadly, were indicative of greater stasis.

To test this “post-hoc prediction”, we added the leadership identification flights (a heterogeneous mix of birds) to the analysis. However our results were that homogenous groups did not show significantly increased stasis, unless homogeneous “follower” phenotypes were removed from the analysis. It seems reasonable that a category as broad as “homogeneity” will not explain the nuances of stability in flock positioning.

Our work thus draws into question the value of being a “follower”. Recent work has suggested that leaders could be exposed to greater risk [81]. Here, by measuring predation events of “virtual prey” (pre-defined in their leadership behaviour) by real fish predators, Ioannou et al. [81] revealed that individuals on the leading edge suffer greater predation. Additionally, a “follower personality” could be maintained by frequency-dependence [46]. Here, follower phenotypes were beneficial in populations with a majority of leader phenotypes in evolutionary simulations, owing to their social-tendency, which kept groups cohesive and thus safer from assumed “threat” [46]. Alternatively, there could be no benefit to followership), with followers making the “best of a bad job” [82], owing to their intrinsic handicap in either genetics or ontogeny.

### BROADER IMPLICATIONS

A question arises that, if flock stasis is so beneficial, why would all individuals/groups not exhibit similar and high levels of stasis? Firstly more experienced individuals may have superior knowledge of a particular route [53,83,84], which may allow for less conflict and thus greater stasis. Conversely, if their “preferred routes” are in conflict, groups of experienced and unexperienced individuals may be less likely to achieve high stasis. Linked to experience is age, whereby developmental progress may be necessary to flock optimally also [84]. Juvenile white ibis (*Eudocimus albus*) and northern bald ibis (*Geronticus eremita*) have both been shown to increase their time participating in optimal V-formation flocking [19, 85] over ontogeny [84, 86]. Whether physiological development and/or cognitive development drives the juveniles’ increase in optimal group-level behaviour, is still unknown. Flock stasis in pigeons could prove a useful model system, owing to the simplicity of experimental manipulations in this system.

It is also important to note that the fitness of individuals will rarely be driven by one facet (e.g. speed/energy optimisation) alone. Instead, the benefits conferred to different individuals/phenotypes will likely be context dependent [28,81,87]. Demonstrating the benefits of an optimal phenotype (or phenotypic composition) in one context could not provide a genuine inference of fitness in the tested animals, whether wild or captive/semi-captive. Nevertheless, in captive animal systems, we can identify a known evolutionary pressure in birds (here, the costs of flight [24]), and test, via manipulations, the influence of individual phenotype, and group phenotypic composition (e.g. [87]). Therefore, we can learn about one important component of life history in isolation, and derive conclusions about individual and group-level success [23].

We provide the first evidence for optimal group compositions in flight, which suggests benefit for individuals which choose when to leave or join groups adaptively [23]. This would require individuals to assess the phenotypic composition of the group before making a decision, based upon the individuals which comprise the group [23]. Such dynamics have been verified experimentally in mixed species flocks, where the removal of a “nuclear” species will dramatically reduce the likelihood of “satellite” species to join groups in foraging activities [88]. There is also evidence that individuals within species can choose to join groups with particular phenotypic compositions. Pruitt and Goodnight [89] found that groups of social spiders *Anelosimus studiosus* re-formed previously optimal phenotypic compositions when experimentally perturbed. This could be explained by simple “emergence”, with individual actions in response to the environment alone resulting in the formation of these groups. Yet this explanation seems less likely than an internal “choice” mechanism, because the composition of *reformed groups* resembled compositions *at their native site*, not the site to which they were experimentally translocated [89]. In light of evidence of “choice” in less cognitively adept spiders [89, 90], it seems plausible that birds could “choose” social partners to benefit from optimal group phenotypic compositions. It is well established that birds can assess mate quality by recognition of phenotype [91, 92]. Whether the energetic costs of flight exert a strong enough selective pressure to favour the evolution of phenotypic recognition related to optimal group composition is likely to be debated, but see [64] for a discussion on how the costs of flight has been – and continues to be – a strong selective pressure, shaping the evolution of birds.

Greater flock stasis is likely beneficial across multiple bird species and flock types, including both cluster flocks and V-formations. Birds which fly in V-formation are – conversely to cluster flocking birds – famous for achieving positive aerodynamic interactions via the wake left behind other birds [20, 79], which is thought to be a property of both the birds’ spatial positioning [18] and phasing of their wing-flaps [19]. Flock stasis relates directly to spatial positioning, and as an easily measurable metric, could be useful in the further study of V-formations. We know that V-formations are rarely optimal in their structure at any given moment in time [18, 19], and that the stability of V-formations can improve through learning [84]. Flock stasis can also be modified for use in V-formations, by, for example, separating the dorso-ventral, and cranio-caudal components of stasis, where both components are known to be of crucial importance [19]. Therefore, in birds which travel in groups, especially when group flight represents a large component of a species’ life history, achieving greater flock stasis through learning or choice of social partners could be adaptive.

Effective use of space can be important in terrestrial animals too, including humans. In human crowd disasters, for example, movement pathways and space use have been identified to influence the chances of stampede [63]. Additionally, turning movements can cause trampling and stampede in humans [93], and ants [94]. Ants are deemed useful in evacuation dynamics due to their evolutionary history of travelling *en-masse* in the same direction [94, 95]. As [94] phrased the idea in their empirical work on turning movements and stampede in ants: “humans have been dealing with traffic congestion for only a few hundreds of years, while ants have been dealing with congestion during millions of years of evolution.” This is also true of birds, where effective collision avoidance could potentially be difference between life and death [33, 45]. Of course, the major caveat to the usefulness of our work in human crowd disasters is that we were looking at relatively “steady” travel. Nevertheless, a holistic framework of effective group travel, identifying differences between non-panicked and panicked groups across taxa could be fruitful.

## CONCLUSION

Flock stasis is a novel metric, which was a consistent predictor of flight speed across all group flights in the study, and robust to a sensitivity analysis of arbitrary parameters. Nevertheless our understanding of how individual phenotype and group phenotypic composition interact to predict flock stasis is not yet conclusive. This inconclusiveness is due to “leader groups” and “homogeneous mass groups”, which, while significant predictors of stasis in initial models, were not entirely robust to sensitivity analyses on arbitrary parameters. Regardless, we introduce a metric, predictive of flight speed at no observable energetic cost, to the combined fields of aerodynamics and group movement worthy of further exploration.

Our conclusions can be applied to wild animal systems and towards a broader understanding of animal ecology. A major current challenge in behavioural biology is to understand how changing environments (i.e. human-driven reduction in naturally occurring resources) may impact social groupings [96] and animal movement ecology [97]. We have evidence that group composition can impact the cost of group flight. If groups need to fly further to reach new resources in the future, are we likely to see a movement towards particular phenotypic compositions, or are the relative benefits of heterogeneity (e.g. [87]) still more powerful? This is a question, which we provide some foundational principles toward, but which will require a collective effort by empiricists (both wild and captive behaviourists) and theoreticians to address. This work also further adds to the body of work demonstrating the opportunities that homing pigeons offer in terms of researching flocking dynamics, personality, energetics and collective behaviour [98–110].

## Acknowledgements.

Funding for this study was provided by the following grants to S.J.P: A Royal Society Research Grant (R10952), a Royal Holloway Research Strategy Fund, and an endowment from the late Professor Percy Butler. We also would like to thank Maté Nagy for help with the analysis of leadership hierarchies; Lucy Taylor for accelerometry R-scripts; Robin freeman for help with GPS waterproofing methods; Linda Garrison and Steven Lang for proof-reading of the introduction; and Emily Shepard, Dora Biro, Liz Greenyer, Francesco Santi, Danai Papageorgiou, and Andrea Perna for useful discussion.

## Notes

### Competing Interest Statement

The authors have declared no competing interest.

